# Active Nucleosome Sliding, Eviction, and Reassembly Dynamics Can Regulate Chromatin Condensate Organization

**DOI:** 10.64898/2026.09.30.755691

**Authors:** Ritu Raj, Ranjith Padinhateeri, P. B. Sunil Kumar

## Abstract

– At the nucleosome scale, chromatin is an actively remodeled polymeric system whose organization is continuously reshaped by ATP-dependent remodelers that slide, evict, and reassemble nucleosomes along DNA. However, how the local structural rearrangements generated by such remodeling events influence condensate-scale organization and dynamics remains poorly understood. Here, we introduce a coarse-grained model of chromatin condensates composed of short polymers that explicitly incorporates enzyme-assisted nucleosome sliding together with nucleosome eviction and reassembly, coupled to Langevin relaxation dynamics. By explicitly accounting for the DNA length sequestered within individual nucleosomes, the model captures the steric constraints and local conformational relaxation that accompany nucleosome sliding, eviction, and reassembly. We find that nucleosome activity drives condensate swelling, enhances condensate-size fluctuations, and promotes internal mixing. In condensates containing both active and passive polymers, differences in nucleosome activity further induce radial segregation, with active polymers enriched near the periphery and passive polymers concentrated in the interior. These results identify active nucleosome remodeling as a physical mechanism for regulating chromatin condensate structure, fluidity, and spatial organization.

## Introduction

Chromatin is a DNA–protein complex that packages genomic DNA inside the eukaryotic nucleus [1]. Its organization emerges across multiple length scales, from chromosome territories and topologically associating domains to local nucleosome-scale structure [2–4]. At the fundamental structural level, chromatin consists of nucleosomes, in which approximately 147 base pairs of DNA are wrapped around a histone octamer, with neighboring nucleosomes connected by linker DNA [5].

Nucleosome organization along DNA is actively regulated by ATP-dependent chromatin remodelers that slide nucleosomes, promote nucleosome eviction, and facilitate reassembly at new locations [6]. Through these processes, remodelers continuously reorganize chromatin structure and modulate DNA accessibility during transcription, replication, and DNA repair [7, 8]. While polymer models have provided important insights into chromatin folding and nonequilibrium dynamics at genomic length scales [9–11], such coarse-grained descriptions typically average over nucleosome-level rearrangements. Conversely, nucleosome-resolved models have primarily focused on passive chromatin organization. Mesoscale studies have shown that linker-DNA length and nucleosome repeat length strongly influence chromatin-fiber compaction and the emergence of distinct higher-order structures [12, 13]. More recent multiscale simulations demonstrated that nucleosome conformational plasticity generates multivalent nucleosome–nucleosome interactions that can promote chromatin condensation [14]. Studies have also connected the multiscale organization of nucleosomes within chromatin condensates to their phase behavior and material properties [15].

In a recent study, we represented remodeler dynamics through an effective interaction-switching model and showed that nonequilibrium remodeling can regulate chromatin condensate organization [16]. However, that approach coarse-grains over nucleosome motion along the polymer backbone and does not explicitly capture the steric constraints and local fiber relaxation associated with nucleosome sliding, eviction, and reassembly.

Here, we address this limitation by explicitly representing nucleosome sliding, eviction, and reassembly along the chromatin fiber, allowing nucleosome positions and occupancy to evolve under local steric and polymeric constraints. We model short chromatin polymers containing nucleosome and linker segments, following *in vitro* chromatin-condensate studies using short nucleosome arrays [17, 18]. Activity is implemented through these stochastic rearrangements along the polymer backbone. By accounting for the DNA length sequestered within nucleosomes, the model captures local fiber relaxation, steric constraints, and transient disruption of nucleosome– nucleosome contacts.

We find that nucleosome activity drives condensate swelling, enhances condensate-size fluctuations, and promotes internal mixing. In condensates containing both active and passive polymers, differences in nucleosome activity further induce radial segregation, with active polymers enriched near the periphery and passive polymers concentrated in the interior. These findings identify local nonequilibrium nucleosome rearrangements as a physical mechanism for regulating chromatin condensate material properties and mesoscale spatial organization.

## Simulation model

Chromatin is modeled as a bead–spring block copolymer composed of compact nucleosome segments connected by semiflexible linker segments (Fig. 1) [20]. DNA wrapping around the histone complex is represented by strong attractive interactions among beads within each nucleosome segment, causing the segment to collapse into a compact structure (Fig. 1(b)). In contrast, linker segments connecting consecutive nucleosome segments along the polymer backbone are composed of self-avoiding beads and remain extended. The system consists of *N*_poly_ = 18 polymers, each containing *N*_*p*_ = 192 beads. Each polymer contains 12 nucleosome segments and 12 linker segments, with nucleosome positions randomly distributed along the polymer. This follows *in vitro* chromatin-condensate studies using ∼12-nucleosome arrays [17, 18]. For *N*_poly_ = 18, the system spans ∼40 kb, comparable to genomic sizes of cellular condensates [19]. A nucleosome segment consists of *N*_*n*_ = 12 beads, whereas a linker segment consists of *N*_*l*_ = 4 beads.

**Fig. 1:**
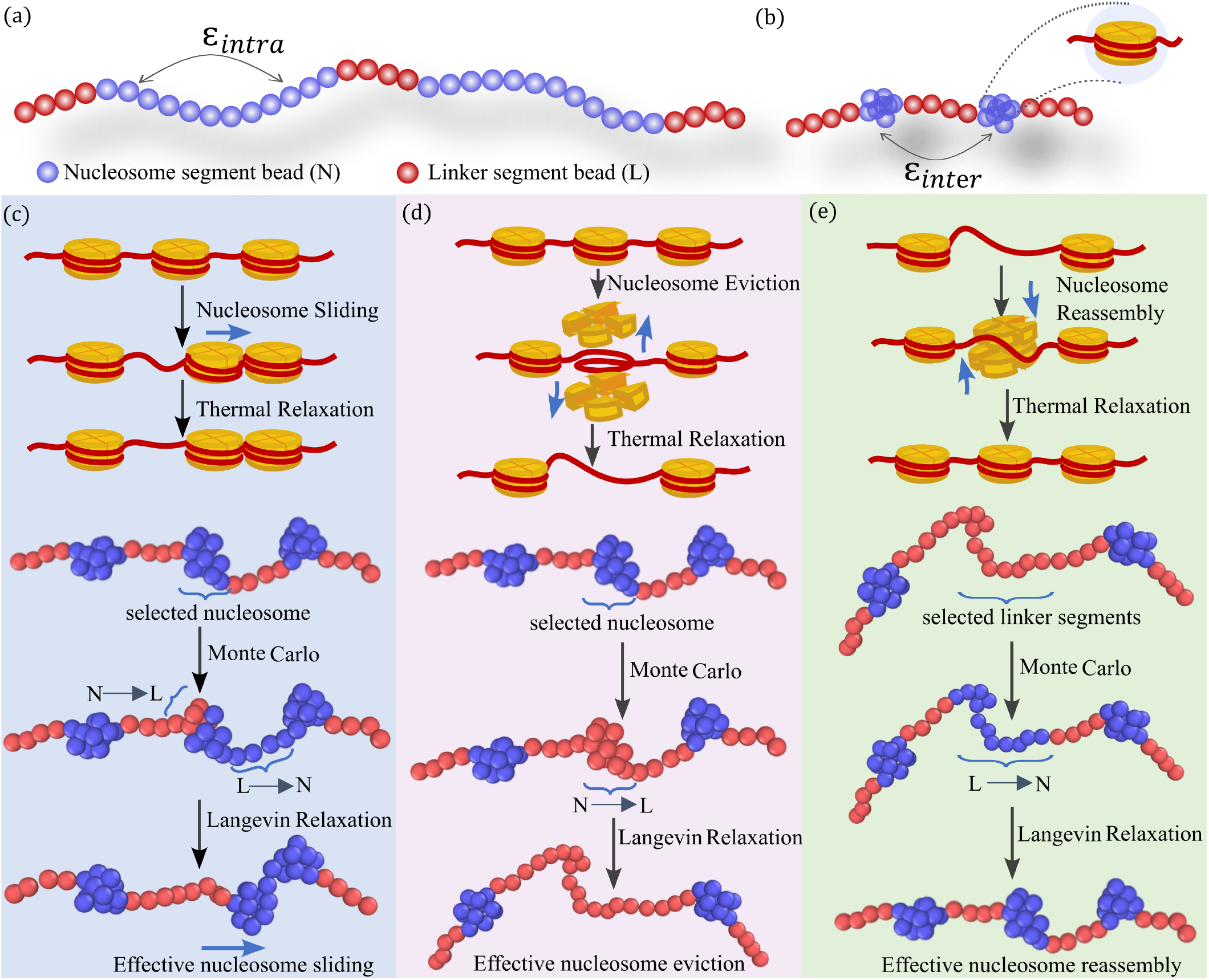
Schematic representation of the coarse-grained chromatin model and nucleosome dynamics. **(a)** Coarsegrained polymer representation of chromatin, consisting of nucleosome and linker segments. Beads within the same nucleosome segment interact via a Lennard-Jones (LJ) interaction with *ϵ*_intra_ = 5*k*_B_*T*, causing the segment to collapse into a compact globule, as shown in **(b)**. The collapsed nucleosome segment represents a nucleosome. Beads belonging to different nucleosome segments interact via an LJ interaction with *ϵ*_inter_ = 0.8*k*_B_*T*. **(c)** *Nucleosome sliding*. A selected nucleosome segment is shifted by *N*_*l*_ beads along the polymer contour by converting the adjacent linker segment into nucleosome beads and the opposite end *N*_*l*_ beads into linker beads. Subsequent LD relaxation for *τ*_LD_ allows the updated segment to collapse, mimicking nucleosome sliding. **(d)** *Nucleosome eviction*. A selected nucleosome segment evicts with probability min(1, *P*_nl_), converting its *N*_*n*_ nucleosome beads into linker beads. During subsequent LD relaxation, the linker region expands, mimicking DNA unwrapping from the histone core and exposing approximately 150 bp of DNA. **(e)** *Nucleosome reassembly*. A linker segment is selected, and if it belongs to a block of three consecutive linker segments, reassembly is attempted with probability min(1, *P*_ln_). Upon acceptance, the three linker segments are converted into one nucleosome segment, which subsequently collapses under the strong intra-nucleosome attraction, *ϵ*_intra_, during LD relaxation, mimicking nucleosome assembly on DNA. *See the Simulation model section for details*.

The position of the *i*-th bead, **r**_*i*_, where *i* = 1, …, *N* (*N* = *N*_poly_ × *N*_*p*_), evolves according to the Langevin equation

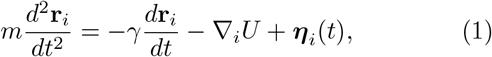

where *γ* is the friction coefficient and ***η***_*i*_(*t*) is a Gaussian random force satisfying ⟨***η***_*i*_(*t*)⟩ = 0, ⟨*η*_*iα*_(*t*)*η*_*jβ*_(*t*^*′*^)⟩= 2*γk*_B_ *Tδ*_*ij*_ *δ*_*αβ*_ *δ*(*t* − *t*^*′*^). Here, *k*_B_ is the Boltzmann constant and *T* is the absolute temperature. Unless stated otherwise, the bead diameter *σ* is used as the unit of length and *k*_B_*T* as the unit of energy. To map the simulation length scale to real units, the average linker length is taken as 50 bp and the DNA bead diameter *σ* is set to 12.5 base pairs (bp). The microscopic time unit is defined as 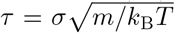, where *m* is the bead mass. The dimensionless friction coefficient is set to 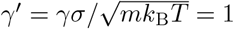.

The total potential energy of the system is given by

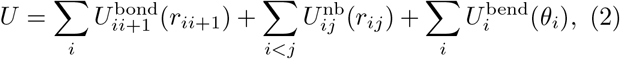

where *r*_*ij*_ = |**r**_*i*_ − **r**_*j*_|. The first term accounts for bonded interactions between consecutive beads along the polymer, the second term accounts for nonbonded interactions, and the third term describes bending rigidity along the polymer.

The bonded interaction is described by the finitely extensible nonlinear elastic (FENE) potential,

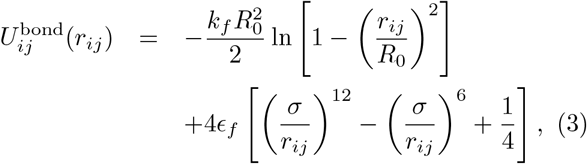

where *k*_*f*_ = 10, *R*_0_ = 1.5, and *ϵ*_*f*_ = 0.305 [20].

Nonbonded interactions between beads in the nucleosome segments, henceforth referred to as nucleosome beads, are modeled using a truncated Lennard–Jones potential,

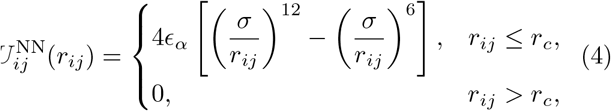

where *r*_*c*_ = 2.5. The interaction strength is set to *ϵ*_*α*_ = *ϵ*_intra_ for bead pairs within the same nucleosome segment and *ϵ*_*α*_ = *ϵ*_inter_ for bead pairs belonging to different nucleosome segments. The value *ϵ*_intra_ = 5 maintains each nucleosome segment in a compact collapsed state, whereas *ϵ*_inter_ = 0.8 provides a weaker attraction between distinct nucleosomes and promotes condensate formation.

Nonbonded interactions of beads in the linker segments, henceforth referred to as linker beads, with all other beads are modeled using the repulsive WCA potential,

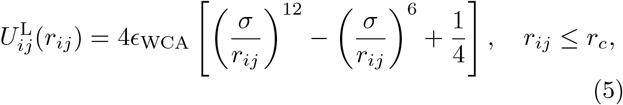

and 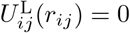 otherwise. Here, *r*_*c*_ = 2^1*/*6^ and *ϵ*_WCA_ = 1. The bending rigidity is introduced through

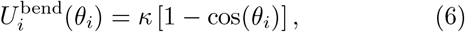

where *θ*_*i*_ is the angle at *i* between consecutive bond vectors. We set *κ* = 10 when the angle is formed by three consecutive linker beads and *κ* = 0 otherwise.

Enzyme-assisted nucleosome dynamics, including sliding, eviction, and reassembly, are incorporated through Monte Carlo (MC) moves coupled to Langevin dynamics (LD) (Fig. 1). After every LD interval of *τ*_LD_ = 50*τ*, an MC step is performed. During each MC step, *N*_*s*_ DNA segments are selected at random. For each selected segment, a nucleosome sliding move is attempted with probability *P*_*s*_, while a nucleosome eviction or reassembly move is attempted with probability *P*_*d*_ = 1 − *P*_*s*_. These moves are implemented by relabeling the coarse-grained beads representing nucleosome and linker DNA; consequently, bead positions remain unchanged during the MC step, and subsequent structural relaxation is governed entirely by the LD evolution.

In a sliding move, a segment is selected at random. If the selected segment is a nucleosome segment, it is slid either forward or backward along the polymer contour, with both directions chosen with equal probability. To perform the slide, the *N*_*l*_ beads of the adjacent linker segment in the chosen direction are converted into nucleosome beads, while *N*_*l*_ beads at the opposite end of the nucleosome segment are converted into linker beads. This effectively translates the nucleosome segment along the polymer contour by a length *N*_*l*_ (Fig. 1(c)). The move is rejected if the selected segment is not a nucleosome segment or if the adjacent segment in the chosen direction is not a linker segment.

In an eviction/reassembly move, a segment is chosen at random. If the selected segment is a nucleosome segment, the *N*_*n*_ beads of the nucleosome are switched into linker beads with probability min(1, *P*_nl_) (Fig. 1(d)). Conversely, if the selected segment is a linker segment that forms part of three consecutive linker segments (since *N*_*n*_ = 3*N*_*l*_), it is switched into a nucleosome segment with probability min(1, *P*_ln_) (Fig. 1(e)).

When accepted, the eviction move yields three consecutive linker segments of size *N*_*l*_, corresponding to ∼150 bp of bare DNA [5], while the reassembly move converts three consecutive linker segments into a nucleosome segment of size *N*_*n*_. The strong intrasegment interaction collapses this segment into a globule, mimicking the wrapping of ∼150 bp of DNA around the histone core [5].

The probability of converting a nucleosome segment into linker segments, corresponding to nucleosome eviction, is given by

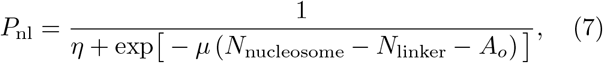

whereas the probability of converting three consecutive linker segments into a nucleosome segment, corresponding to nucleosome reassembly, is

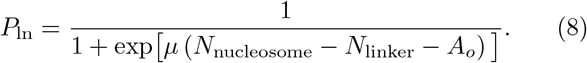

Here, *N*_nucleosome_ and *N*_linker_ are the instantaneous numbers of nucleosome and linker beads, respectively, and 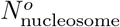 and 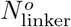 are their steady-state values. The parameter 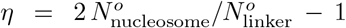 ensures that *N*_nucleosome_ and *N*_linker_ approach their steady-state values. The parameter *µ* controls the magnitude of fluctuations in nucleosome and linker numbers, while 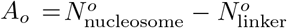 is the asymmetry parameter. These switching probabilities depend on the instantaneous numbers of nucleosome and linker beads and maintain a steady state [20, 21].

In our simulations, the total number of beads is *N* = 3456, with 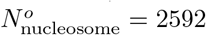 and 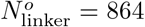, yielding *η* = 5 and *A*_*o*_ = 728. To maintain approximately constant nucleosome and linker numbers with minimal fluctuations, we set *µ* = 1.

At the end of every MC move, during which the relabeling associated with a sliding, eviction, or reassembly event is completed, the system is evolved by Langevin dynamics for *τ*_LD_ before the next MC attempt is made (Fig. 1). During this relaxation period, newly formed nucleosome segments collapse into compact nucleosome structures, while newly formed linker segments relax as extended solvophilic segments. We choose *τ*_LD_ = 50*τ*, which is the time required for an initially straight nucleosome segment to collapse into its compact structure after relabeling. This choice sets the characteristic timescale for completing a nucleosome sliding, eviction, or reassembly event.

The *activity, A*, is defined as the fraction of segments selected for Monte Carlo moves during each relaxation in-

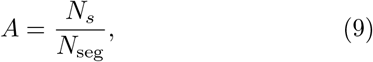

where *N*_*s*_ is the number of segments selected for MC moves and *N*_seg_ is the total number of segments. Thus, increasing *A* increases the frequency of nucleosome sliding, eviction, and reassembly events. Since a Monte Carlo step is initiated after every LD interval of *τ*_LD_, *N*_*s*_ sets the activity timescale, defined as the LD interval required for every segment to be picked once, on average, in the MC move.

## Results and discussion

### Formation of chromatin condensate

A condensate of 18 chromatin polymers, each containing 12 nucleosomes, was simulated. Simulations were initialized from straight polymer conformations, and attractive internucleosome interactions drove the coalescence of the polymers into a single condensate. The system was evolved until it reached a steady state.

Nucleosome activity (sliding, eviction, and reassembly) was introduced only after condensate formation. When activity is introduced before condensate formation, frequent sliding, eviction, and reassembly events disrupt the formation of stable nucleosome–nucleosome contacts required for condensate formation, and we do not observe condensate formation within our simulation timescale. This effect is particularly strong when the activity timescale, *τ*_LD_/*A*, is shorter than the polymer relaxation time, *τ*_*R*_ ≈ 3.22 × 10^4^*τ*, estimated from the 1/e decay of the normalized end-to-end vector autocorrelation function, 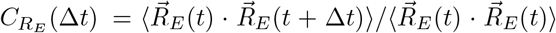. Under these conditions, chromatin polymers do not have sufficient time to establish persistent internucleosome contacts, and stable condensates do not form. Therefore, in all subsequent simulations, nucleosome activity was introduced only after the passive condensate had reached steady state. For activity values higher than *A* = 0.4, polymers escape from the condensate, preventing the formation of a stable condensate. Therefore, we restrict our analysis to *A* ≤ 0.4.

Thus, our protocol probes the response of a stable chromatin condensate to ongoing nucleosome activity, rather than the kinetics of condensate assembly. This is consistent with *in vitro* experiments showing that remodelers present before condensation can inhibit condensate formation [18].

### Effect of nucleosome activity on chromatin condensates

We consider the case where eviction/reassembly and sliding attempts are performed with equal probability (*P*_*d*_ = *P*_*s*_ = 0.5). Upon switching on activity in a preformed steady-state condensate, we observe gradual swelling of the condensate. This swelling is quantified by the condensate radius of gyration, R_*g*_(*t*), defined as [22]

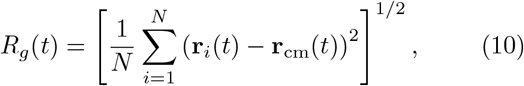

where *N* is the total number of beads in the condensate and **r**_cm_(*t*) is the condensate center of mass. As shown in the inset of Fig. 2(a), R_*g*_(*t*) initially increases with time and subsequently saturates to a steady-state value, ⟨R_*g*_⟩. The steady-state radius of gyration increases with activity, indicating activity-induced swelling of the chromatin condensate, as shown in Fig. 2(a). Representative snapshots are shown in Supplementary Material Fig. S1. A similar trend is observed for *N*_poly_ = 27 in Fig. S2.

**Fig. 2:**
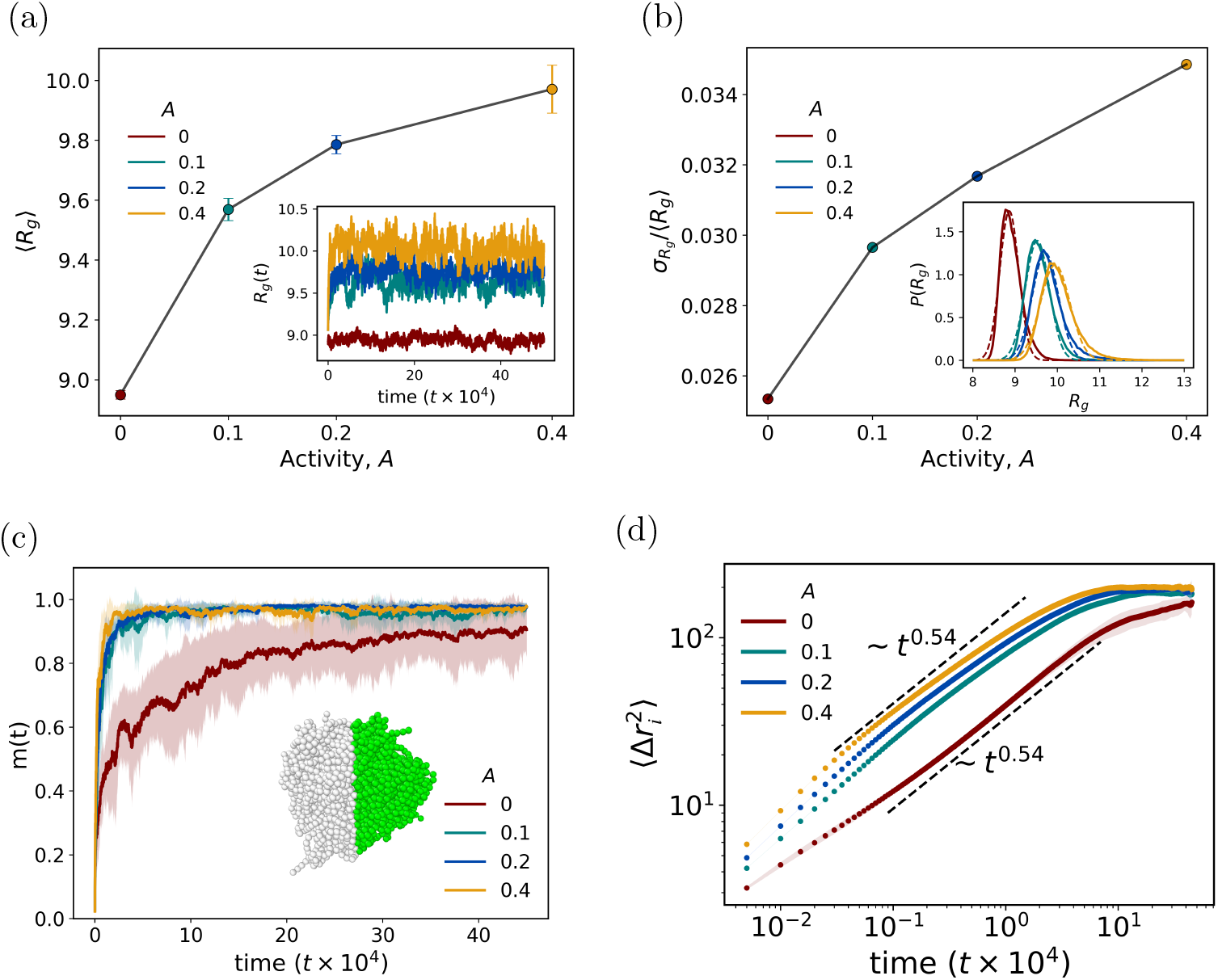
**(a)** Average condensate radius of gyration, ⟨*R*_*g*_ ⟩, as a function of activity, *A*. The inset shows the time evolution of *R*_*g*_ (*t*) after activity is switched on in a preformed condensate. **(b)** Normalized fluctuation of the condensate radius of gyration, 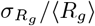, as a function of activity, obtained from the standard deviation of the steady-state *R*_*g*_ distribution. The inset shows the corresponding *R*_*g*_ distributions, with dashed lines denoting Gaussian fits. **(c)** Monomer mixing parameter, *m*(*t*), for passive and active condensates. The inset illustrates the initial labeling used to compute the mixing parameter. **(d)** Monomer mean-square displacement (MSD) in the condensate center-of-mass frame, 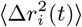, for different activities. Shaded regions in (c) and (d) denote the standard deviation.

The observed swelling arises from a competition between attractive nucleosome–nucleosome interactions that favor condensation and the configurational entropy generated by active nucleosome rearrangements. Nucleosome sliding, reassembly, and eviction continuously remodel intracondensate contacts, enabling polymers to sample increasingly expanded conformations. As activity increases, this entropic contribution progressively counteracts the attractive interactions, resulting in a larger condensate radius of gyration. When the segment-switching timescale becomes shorter than the polymer relaxation time, *τ*_*R*_, polymers cannot fully relax into compact configurations before further rearrangements occur, amplifying the swelling.

In addition to increasing the steady-state average condensate size, activity also enhances fluctuations in the condensate radius of gyration, as shown in Fig. 2(b) (See also Supplementary Movie S1). We quantify these fluctuations by the normalized standard deviation, 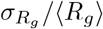, obtained from Gaussian fits to the steady-state R_*g*_ distributions shown in the inset of Fig. 2(b). The normalized fluctuation increases monotonically with activity, indicating that higher nucleosome activity not only swells the condensate but also makes its size more dynamically fluctuating.

To examine whether activity-induced swelling is accompanied by enhanced fluidity of chromatin polymers, we computed the monomer mixing parameter, *m*(*t*). To calculate *m*(*t*), monomers in one half of the condensate are initially labeled as type A, while monomers in the other half are labeled as type B, as illustrated in the inset of Fig. 2(c). At each time, we compute pairwise distances between all monomers, *d*_all_, and between type-A and type-B monomers, *d*_cross_. These distances are then binned to obtain the corresponding distance distributions, *P*_all_(*r, t*) and *P*_cross_(*r, t*). The monomer mixing parameter is defined as *m*(*t*) = 1 − *M*(*t*), where

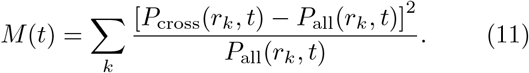

Here, *k* indexes the distance bins and *r*_*k*_ denotes the center of the *k*-th bin. The quantity *M*(*t*) is normalized to lie between 0 and 1; therefore, *m*(*t*) = 0 corresponds to an unmixed state, whereas *m*(*t*) = 1 indicates complete mixing [16, 23].

In the presence of activity, monomers intermix substantially faster than in the passive condensate, as shown in Fig. 2(c). We supplement this analysis by calculating the enhanced monomer mobility using the monomer mean-square displacement (MSD) in the condensate center-of-mass frame, 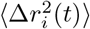. As shown in Fig. 2(d), for both passive and active condensates, the MSD remains subdiffusive, with an approximate scaling of ∼*t*^0.54^, consistent with Rouse-like dynamics [24]. Increasing activity does not significantly change the scaling exponent but shifts the MSD curves upward, indicating enhanced effective monomer mobility. Thus, nucleosome activity promotes faster internal rearrangement and monomer mixing while preserving Rouse-like dynamics by increasing the effective temperature.

### Condensates of polymers with different nucleosome spacing and activity

We next consider mixed condensates containing equal numbers of passive and active polymers. Nucleosome activity is switched on only for half of the polymers, with the activity parameter fixed at *A* = 0.1. For the passive polymers, we examine two nucleosome-spacing scenarios: (i) uniformly spaced nucleosomes, in which neighboring nucleosome segments are separated by one linker segment, and (ii) randomly spaced nucleosomes, in which nucleosome segments are randomly distributed along each polymer while keeping the total number of nucleosomes fixed at 12 per polymer.

To quantify the spatial organization of active and passive polymer populations within the condensate, we computed the radial number density of polymer beads,

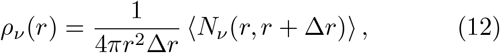

where *v* denotes the active or passive polymer population, and *N*_*v*_ (*r, r* + Δ*r*) is the number of beads belonging to population *v* located in a spherical shell between *r* and *r* + Δ*r* from the condensate center of mass.

The radial bead density profiles reveal clear segregation between active and passive polymers (Fig. 3(a,b)). Passive polymers preferentially occupy the condensate interior, whereas active polymers are enriched near the periphery. This segregation originates from the continual remodeling of nucleosome positions along active polymers. Repeated nucleosome sliding, eviction, and reassembly generate a heterogeneous nucleosome-spacing pattern with a broad distribution of linker lengths, including longer linker segments. As shown earlier, such heterogeneous spacing favors a more expanded, less densely packed polymer conformation [16]. Consequently, in mixed condensates, the more densely packed passive polymers preferentially occupy the core, while the lower-density active polymers are displaced toward the interface, giving rise to the observed core–shell organization.

**Fig. 3:**
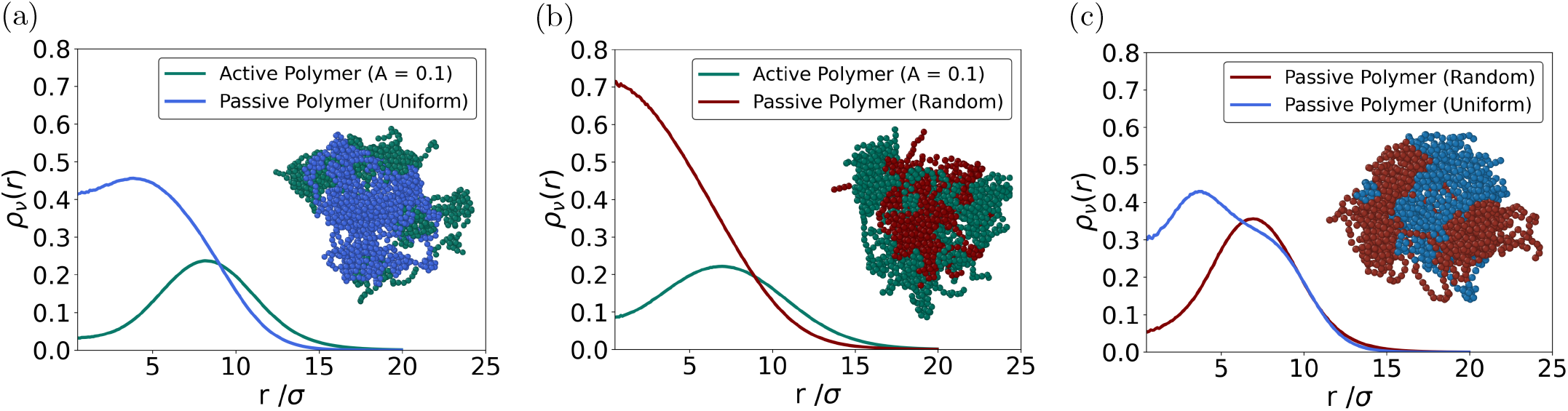
Radial density profiles, *ρ*_*v*_ (*r*), of polymer beads in chromatin condensates with different nucleosome-spacing arrangements and activity. **(a)** Equal mixture of active and passive polymers with uniformly spaced nucleosomes. **(b)** Equal mixture of active and passive polymers with randomly spaced nucleosomes. **(c)** Passive condensate containing an equal mixture of polymers with randomly and uniformly spaced nucleosomes. Insets show representative cross-sectional cuts of the condensates, including fiber beads satisfying |*z* − *z*_COM_| *<* 2*σ*. Colors identify the corresponding fiber types as indicated in the legends.

To test this interpretation, we compare the results with simulations of mixed condensate containing equal numbers of polymers with uniform and random nucleosome spacing in the absence of activity [16]. The radial density profiles show similar segregation, with uniformly spaced polymers preferentially occupying the condensate interior and randomly spaced polymers enriched toward the outer region, as shown in Fig. 3(c). This supports the idea that activity-induced heterogeneity in nucleosome spacing is sufficient to induce the radial organization observed in active–passive condensates. However, the distributions obtained for passive and active mixtures are distinctly different.

For mixed condensates containing active polymers and passive polymers with random nucleosome spacing, active polymers preferentially occupy the outer region. This occurs because activity continuously disrupts and rearranges nucleosome–nucleosome contacts. In contrast, randomly spaced passive polymers, despite their heterogeneous linker lengths, retain stable nucleosome–nucleosome contacts and therefore remain denser and enriched in the condensate interior.

## Conclusion

Overall, our results show that local nonequilibrium nucleosome rearrangements can regulate condensate swelling, internal mixing, and spatial organization. The model provides a minimal physical framework for connecting enzyme-assisted nucleosome dynamics to mesoscale chromatin organization. The activity-induced condensate swelling reported here is consistent with *in vitro* observations [18]. Enhanced internal mixing could be tested using Fluorescence Recovery After Photobleaching, while the predicted radial segregation of active and passive chromatin fibers could be examined by cryoelectron microscopy. The qualitative agreement between the present explicit sliding–eviction–reassembly model and earlier effective interaction-switching descriptions suggests that these activity-induced changes are robust outcomes of local nonequilibrium remodeling [16]. Future extensions incorporating explicit remodeler proteins, histone-tail interactions, electrostatics, and hydrodynamic effects could further bridge this framework with *in vivo* chromatin organization.

Computational support from the IIT Madras AQUA Super Cluster Facility is gratefully acknowledged. PBSK acknowledges financial support from ANRF, India, under the J. C. Bose Grant (ANRF/JBG/2025/000187/PS).

