## Supplementary Material for "Active Nucleosome Sliding, Eviction, and Reassembly Dynamics Can Regulate Chromatin Condensate Organization"

In this Supplementary Information, we provide additional figures and movie that complement the results presented in the main text.

### Supplementary Figures

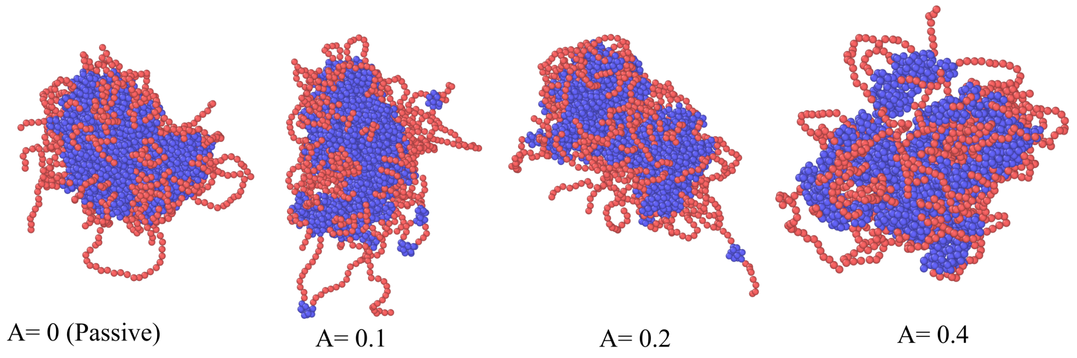


Figure S1: Representative snapshots of chromatin condensates at activities $A = 0$ (passive), 0.1, 0.2, and 0.4. Blue beads represent nucleosome-segment beads, while red beads represent linker-DNA beads.

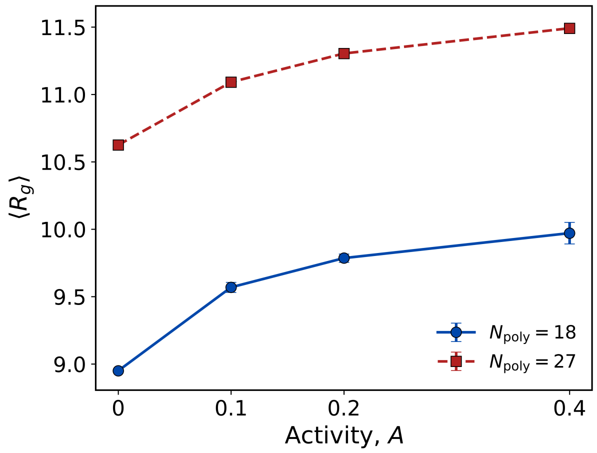


Figure S2: Steady-state average radius of gyration, $\langle R_g \rangle$, as a function of activity for a condensate containing $N_{\text{poly}} = 27$ polymers. Error bars denote the standard deviation.

### Supplementary Movie

**Movie S1. Dynamics of Passive and Active Chromatin Condensates.** Representative movies of chromatin condensates for the passive case, $A = 0$, and the active case, $A = 0.4$.
